# Discovering dynamic plant enzyme complexes in yeast for novel alkaloid pathway identification from a medicinal plant kratom

**DOI:** 10.1101/2023.01.16.524293

**Authors:** Yinan Wu, Chang Liu, Franklin L. Gong, Sijin Li

**Author notes:** Corresponding author: Li, Sijin. These authors contributed equally to this work (Yinan Wu, Chang Liu).

## Abstract

Discovering natural product biosynthetic pathways from medicinal plants is challenging and laborious, largely due to the complexity of the transcriptomics-driven pathway prediction process. Here we developed a novel approach that captures the protein-level connections between enzymes for pathway discovery with improved accuracy. We proved that heterologous protein-protein interaction screening in yeast enabled the efficient discovery of both dynamic plant enzyme complexes and the pathways they organize. This approach discovered complexes and pathways in the monoterpene indole alkaloid metabolism of a medicinal plant, kratom with high success rate. Screening using a strictosidine β-D-glucosidase (MsSGD1) against 19 medium-chain dehydrogenase/reductases (MsMDRs) identified five MsSGD1-MsMDR complexes. Three out of the five interacting MsMDRs were then proven functional, while the remaining 14 non-interacting candidates did not show obvious activities. The work discovered three branched pathways by combining transcriptomics, metabolomics, and heterologous PPI screening and demonstrated a new plant pathway discovery strategy.

---

Plant natural products (PNPs) synthesized by medicinal plants occupy unique structural space and thereby important therapeutic niches. Discovering their biosynthetic pathways enables the biomanufacturing of pharmaceutical PNPs and provides new biochemical knowledge for drug discovery and development. Transcriptomics analysis has proven a powerful method to predict putative PNP pathways by their expression or co-expression “patterns” and led to the elucidation of pathways from multiple medicinal plants, including mayapple^1^, kava^2^, and *Gloriosa superba*^3^. These RNA-level patterns are usually coupled with plant metabolomics analysis, which links the gene expression levels to metabolite concentrations in different plant samples^4^ for enhanced prediction success rate. However, the synthesis of PNPs is a highly regulated and pluri-organelle process^5^, which current multi-omics methods have not fully pictured. The vast majority of genes predicted by transcriptomics analysis are usually found to be irrelevant, despite their expression patterns being highly similar to the characterized pathway genes. The low efficiency of transcriptomics-driven pathway prediction and, consequently, laborious prioritization and characterization efforts afterward^6^ significantly lower the speed of PNP pathway discovery from medicinal plants.

A protein-level pattern that can distinguish pathway enzymes from irrelevant plant proteins precisely might complement the prevalent RNA-centered pathway prediction strategy and provide new mechanistic insights to discover novel PNP pathways. We hypothesize that the well-documented spatial organization machinery in plants, namely dynamic plant enzyme complexes (enzyme complexes for short), can guide pathway discovery based on their unique biochemical and biophysical properties. First, all enzyme complexes discovered to date are solely composed of sequential enzymes in a biosynthetic pathway without any irrelevant proteins^7^. Second, the protein-protein interactions (PPIs) between enzymes in the same complex are highly specific. For example, among the 11 chalcone reductase paralogs in soybean, only one leads to a functional enzyme that interacts with the scaffold enzyme to form the isoflavone enzyme complex^8^. Although not all pathways are organized in the form of complexes, the extensive existence of enzyme complexes in diverse plants and metabolisms opens the door to a large number of pathways that used to be inaccessible using traditional approaches. As plant PPIs are challenging to validate in the native host efficiently, we developed a heterologous PPI screening method in *Saccharomyces cerevisiae* to discover novel enzyme complexes and, correspondingly, the pathways they organize.

We chose a valuable yet less-investigated medicinal plant named kratom^9,10^ (*Mitragyna speciosa*. **Fig. 1a**) to validate our hypothesis. *M. speciosa* produces various monoterpene indole alkaloids (MIAs) with great pharmaceutical potential as opioid agonists^11^ and biased analgesics^12^. However, studies on *M. speciosa* MIA biosynthetic pathways are minimal. The first peer-reviewed draft Kratom genome was just published in 2021^13^. Although no MIA biosynthetic pathways or related complexes were reported from *M. speciosa*, the discovery of multiple enzyme complexes with conserved structures in the well-studied MIA producer, *Catharanthus roseus*^14,15^, provided validated examples to guide our discovery. The nucleus-localized *C. roseus* strictosidine β-D-glucosidase (CrSGD) can interact with three different medium-chain dehydrogenase/reductase (CrMDRs) (**Fig. 1b**), including tetrahydroalstonine synthases 1 and 2 (THAS1 and 2) and heteroyohimbine synthase (HYS), which has been proven by fluorescent fusion protein expression and bimolecular fluorescence complementation (BiFC) assays in *C. roseus* cells^14,15^. In these MIA pathways, CrSGD converts the common MIA precursor, strictosidine, to unstable and cytotoxic strictosidine aglycone, which quickly rearranges to multiple reactive isomers that are immediately converted by CrMDRs to more stable downstream MIAs^16^. Therefore, the interaction between SGD and MDRs is likely related to the biochemical activities of these two types of enzymes and could be common and conserved machinery across various MIA pathways^5^ and species. We postulate that the SGD-MDR interactions could be screened and combined with the established multi-omics strategy to discover novel MDRs and corresponding pathways in *M. speciosa*.

**Fig. 1.**
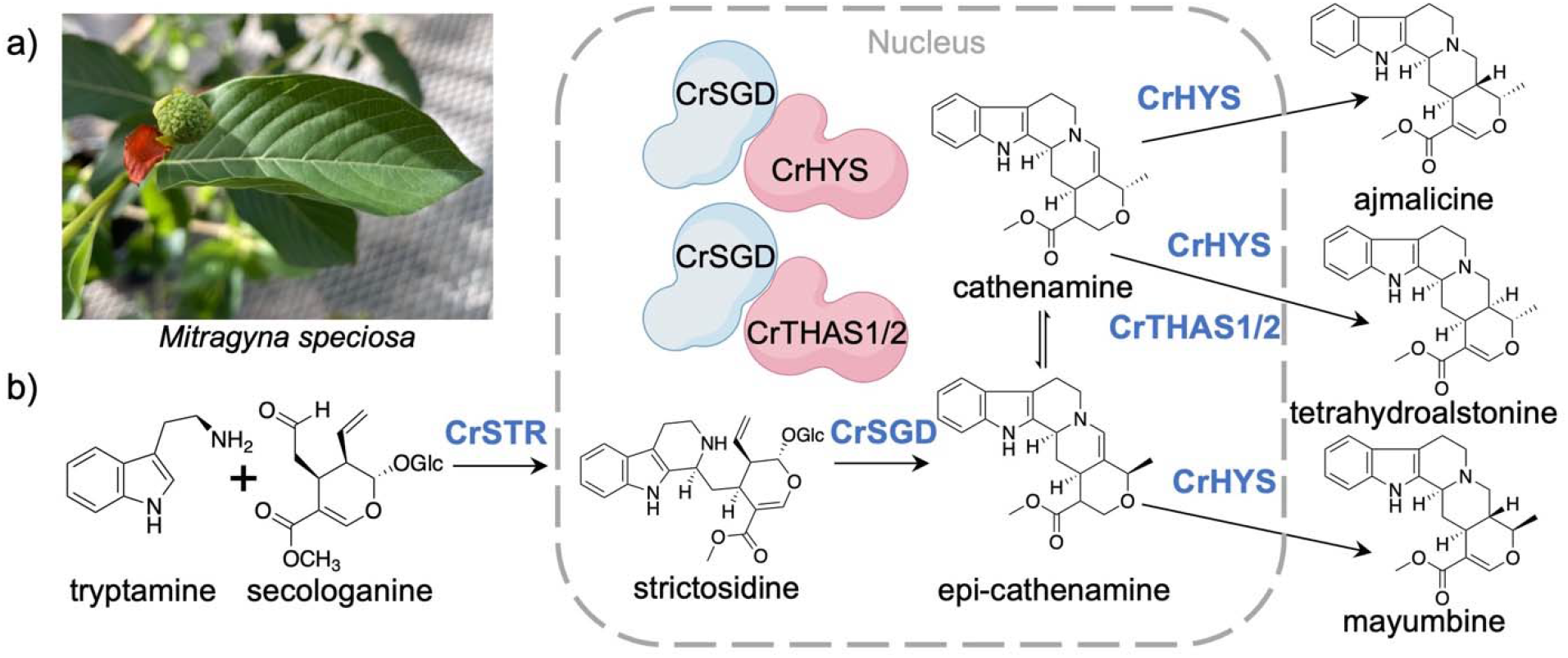
Characterized *C. roseus* MIA pathways and enzyme complexes and *M. speciosa*. a) A photo of *M. speciosa* leaf and flower. b) Characterized *C. roseus* MIA pathways involving enzyme complexes of CrSGD and CrMDRs.

Here we report the discovery of six functional MIA pathway genes and three pathway branches from *M. speciosa* using an integrated approach combining transcriptomics, metabolomics, and the new yeast-based PPI screening method. The multi-omics strategy identified a functional strictosidine synthase (MsSTR) and two functional enzymes showing strictosidine β-D-glucosidase activity (MsSGD1 and MsSGD2), along with 19 prioritized, uncharacterized MsMDRs. Complementary PPI screening methods at the molecular and organelle levels using BiFC and fluorescent protein co-localization assay discovered five MsSGD1-MsMDR complexes. Biochemical characterization further confirmed that three out of the five interacting MsMDRs are functional enzymes converting strictosidine aglycone to downstream MIAs, yielding a 60% success rate in pathway gene prediction. Meanwhile, no products were observed from the remainder 14 MsMDR candidates that did not show apparent signs indicating the formation of complexes. Taken together, the high success rate using PPI screening to discover novel MsMDRs proved that enzyme complex is a promising protein-level pattern for PNP pathway discovery, which has yet to be fully exploited. In summary, our work demonstrated a strategy that uses yeast-based PPI screening for efficient PNP pathway discovery from medicinal plants. This strategy provides an accurate method to complement the prevalent transcriptomics-driven approach for accelerated PNP pathway discovery.

## Results

### Development of yeast-based PPI identification methods to validate the SGD-MDR interaction heterologously

The PPIs between SGD and MDRs have been observed in *C. roseus* cells^14,15^. However, the *in-planta* PPI identification methods are limited by the low throughput and long turnaround time and have yet to be developed in most plants. We developed three methods to identify the PPIs between plant enzymes at the molecular and organelle levels heterologously within the yeast cell, including yeast 2-hybrid (Y2H) assay, yeast-based BiFC, and protein localization analysis using fluorescent fusion proteins. The reported PPI between CrSGD and CrHYS was used to validate the methods. As the yeast strain co-expressing CrSGD and CrHYS could not grow in the Gal4-based Y2H assay^31^, which is possibly due to the SGD-MDR interaction incorrectly blocking the activation of the downstream selective gene, we chose BiFC and protein localization analysis for the following method development.

In the BiFC assay, CrSGD and CrHYS were fused with split mVenus fragments (NV and CV)^17^ in yeast. CV was fused to the N-terminus of CrSGD (CV-CrSGD) to maintain the C-terminal nuclear localization signal (NLS)^18^. NV was fused to the N-termini of CrHYS (NV-CrHYS) and a *Papaver somniferum* methyltransferase, Ps4’OMT^19^ (NV-Ps4’OMT), which is unlikely to interact with *C. roseus* enzymes, as the negative control. Engineered yeasts were cultured for three days and analyzed using a confocal microscope. BiFC results showed that CrSGD and CrHYS interacted in the nucleus (**Fig. 2a**), consistent with the BiFC result in *C. roseus*^14^. No interaction was observed between CrSGD and Ps4’OMT (**Fig. 2b**).

**Fig. 2.**
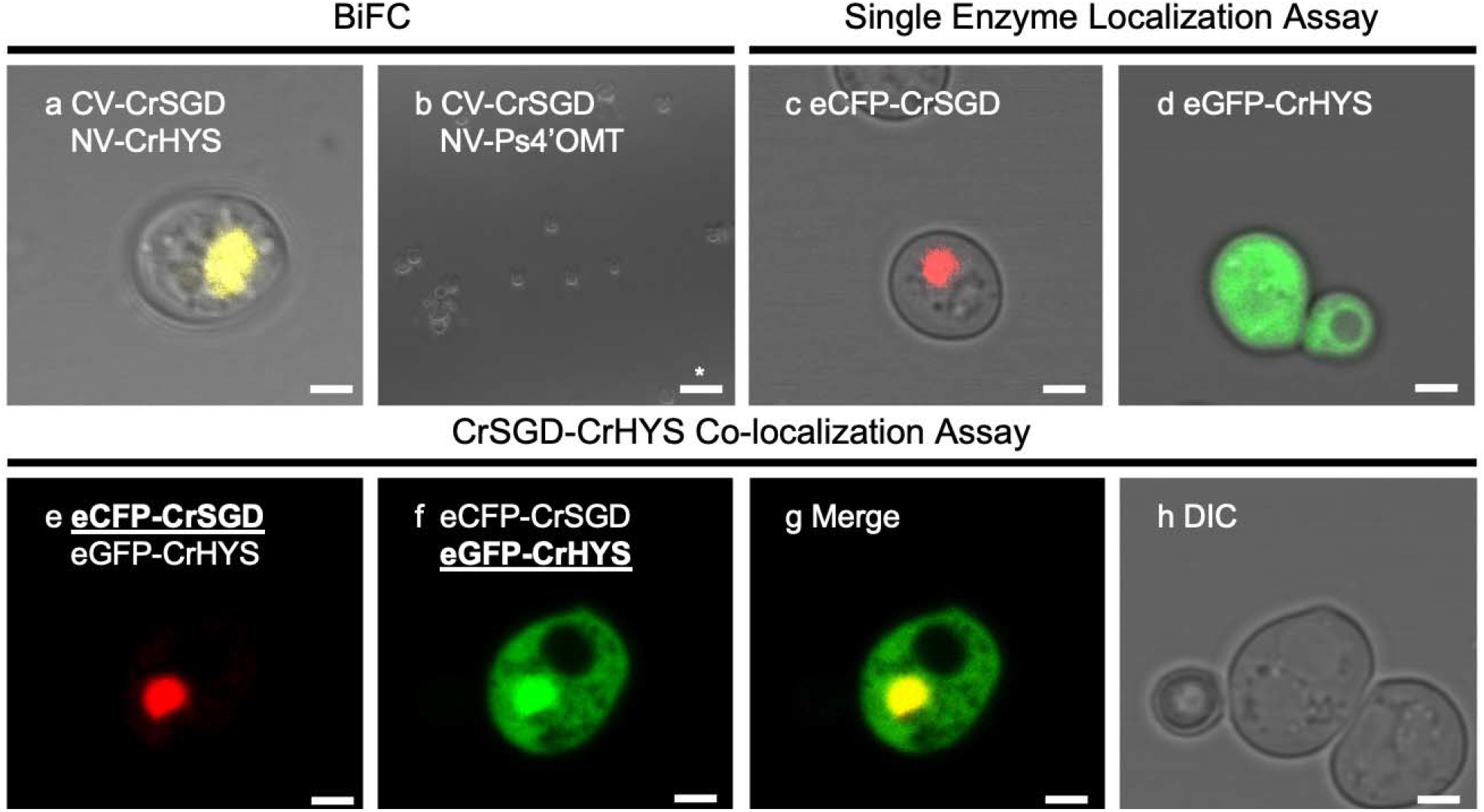
CrSGD-CrHYS interaction validated in yeast using BiFC (a-b) and co-localization analysis (c-h). a-b). CrSGD-CrHYS interaction was analyzed by BiFC in yeast cells co-expressing CV-CrSGD and NV-CrHYS or NV-Ps4’OMT. Fluorescence signal in yellow false color. c-d). Single enzyme localization in yeast cells expressing eCFP-CrSGD (nucleus) or eGFP-CrHYS (cytoplasm), respectively. Fluorescence signals in red or green false colors. e-f). Localizations of co-expressed CrSGD (nucleus) and CrHYS (both nucleus and cytoplasm), respectively. g). Merged localization of co-expressed CrSGD and CrHYS in yellow. h). Cell morphology is observed with differential interference contrast (DIC). All scale bars are 2 μm except for the one labeled with an asterisk (15 μm).

We used the fusion protein co-localization assay to prove the PPI by the subcellular co-localization of CrSGD and CrHYS in yeast. Three yeasts expressing eCFP-CrSGD, eGFP-CrHYS, or both were examined under a confocal microscope. Our results showed that CrSGD was always targeted to the nucleus when expressed individually **(Fig. 2c)** or co-expressed with CrHYS (**Fig. 2e**), as reported *in planta*^18^. In the yeast expressing CrHYS alone, CrHYS localized mainly in the cytoplasm of yeast **(Fig. 2d)**, while it localized in both the nucleus and cytoplasm of *C. roseus* and showed a preferentially nuclear location^14^. Co-expressing CrHYS with CrSGD relocalized a large portion of CrHYS to the nucleus, similar as observed in *C. roseus* (**Figs. 2f and 2g**) and showed a nucleocytoplasmic localization. We speculate that single CrHYS was not targeted to the yeast nucleus because of the uncommon NLS of CrHYS, a plant-specific KKKR sequence that might not be recognized effectively in yeast^20^. The cytoplasmic localization of CrHYS in the absence of CrSGD and, correspondingly, the re-localization of CrHYS with CrSGD into yeast nucleus, provided more convincing evidence that the assembly of the CrSGD-CrHYS complex is driven by molecular-level PPI rather than simple protein compartmentalization. In contrast, when CrHYS was naturally targeted to the plant nucleus, it was difficult to assert that a complex was indeed assembled *in planta* as both CrSGD and CrHYS would be targeted in the nucleus of *C. roseus* independently.

### Metabolomics analysis and *de novo* transcriptome assembly of *M. speciosa*

Previous studies and our metabolomics analysis of different *M. speciosa* tissues (mature leaf, young leaf, root, and flower) have proven that the leaf of *M. speciosa* is rich in MIAs, e.g., strictosidine and mitragynine. The notable differences in MIA accumulation between mature and young leaf tissues indicated different transcript accumulation profiles for the MIA biosynthetic genes. Therefore, we started with the *de novo* assembly of the *M. speciosa* leaf transcriptome using mature (M) and young (Y) leaf samples from two types of *M. speciosa* plants, “Rifat Thai” (T) and “Malaysian” (M). Four types of leaf samples (T_M, T_Y, M_M, and M_Y) led to a transcriptome with 262783 Trinity transcripts and 129489 putative complete or incomplete open reading.

### Identification of one functional MsSTR, two functional MsSGDs, and 19 MsMDR candidates

We started with gene mining to identify MsSTR as the first enzyme in the MIA biosynthetic pathway using the characterized STRs from *C. roseus* and *Rauvolfia serpentina* (CrSTR and RsSTR)^5^ against the *M. speciosa* transcriptome. The MsSTR candidate was functionally characterized in yeast fed with secologanin and tryptamine and was compared with a functional CrSTR variant (trCrSTR)^21,22^ due to the lack of strictosidine standards. Liquid chromatography-mass spectrometry (LC-MS) results revealed that both MsSTR and trCrSTR produced the same product that showed the identical mass/charge ratio as strictosidine (*m/z* 531.2337, 20 ppm), which was not observed in the negative controls (cultural medium without yeast, and with yeast harboring a blank vector) (**Fig. 3a**). The concentrations were estimated to be 0.04 and 0.10 mM (**Fig. 3b**), respectively, according to the consumption of tryptamine. Sequence analysis showed that MsSTR has an N-terminal vacuole signal peptide similar to CrSTR and RsSTR^18^, as well as 15 conserved residues as CrSTR and RsSTR, which have proven critical in the substrate binding process^23,24^.

**Fig. 3.**
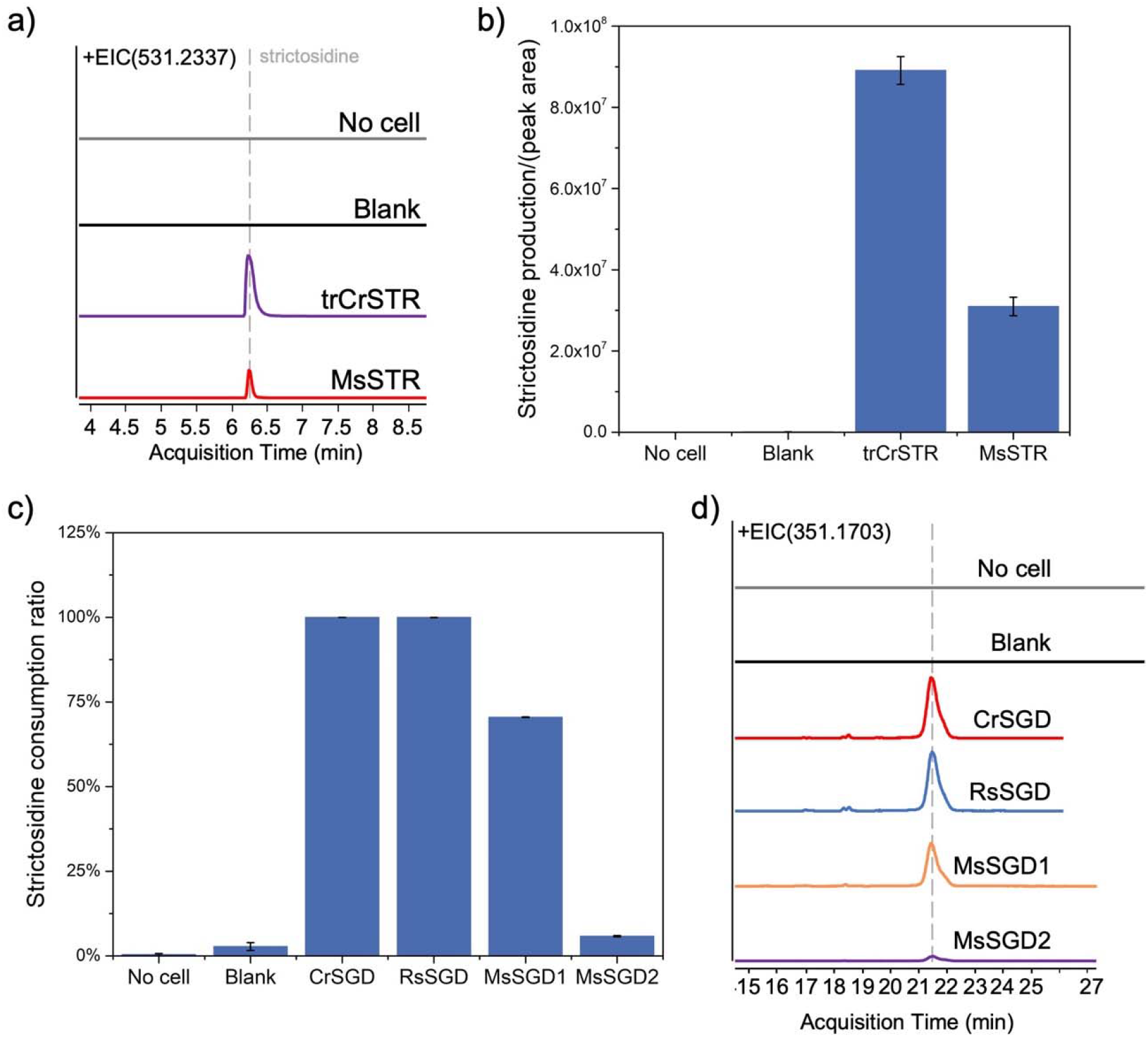
Biochemical characterization of MsSTR and MsSGDs. a) Extracted ion chromatogram (EIC) of strictosidine (*m/z* 531.2337, 20 ppm) after 48-hour fermentation. b) Integrated peak areas of strictosidine produced in different constructs. c) Strictosidine consumption by SGDs in cell lysate assays. d) EIC of *m/z* 351.1703 in cell lysate assays. Bars and error bars represent mean□± □s.d. (n□=□3).

Gene mining using CrSGD and RsSGD identified MsSGD1 and MsSGD2, which are supposed to convert strictosidine to strictosidine aglycones and to interact with downstream MsMDRs. MsSGD1 and MsSGD2 share 61% sequence similarity. Engineered yeasts that expressed the four SGDs and a blank control were lysed and tested with ∼0.1 mM of crude strictosidine (prepared from trCrSTR test) at 30□ for one hour for characterization. MsSGD1 and MsSGD2 consumed 70% and 6% of the strictosidine, while CrSGD and RsSGD consumed 99% and 98% (**Fig. 3c**). All SGDs produced the same products (*m/z* 351.1703 and *m/z* 369.1809, 20 ppm) (**Fig. 3d**), which showed the same m/z as reported strictosidine aglycone isomers^16,25^. Both MsSGD1 and MsSGD2 have the plant β-glucosidases conserved sequence^26^ and four identical active residues with CrSGD and RsSGD^27^. Similar to the C-terminal NLS reported in CrSGD and RsSGD^18^, there are also C-terminal bipartite NLS (522-**<u>KR</u>**TLEDHEDFVS**<u>KKR</u>**<u>L</u>**<u>R</u>**Q-539) and (519-**<u>KR</u>**ALSNGDLEANSNVEEIP**<u>KKK</u>**<u>VL</u>**<u>K</u>**F-544) in MsSGD1 and MsSGD2. MsSGD1-2 transcripts accumulate high in the mature leaves, while MsSTR accumulates preferentially in the young leaves. The opposite expression profiles are likely related to the higher accumulations of strictosidine in the young leaves of *M. speciosa*, which was also demonstrated in *C. roseus*^18^.

We then searched for putative MsMDRs, which are potential MIA pathway enzymes downstream of MsSGD and might interact with SGD to form complexes in *M. speciosa*. 190 Trinity “genes” were functionally annotated as MDRs or functionally similar enzymes (e.g., alcohol dehydrogenases) in the transcriptome. 19 putative *MsMDR* genes were selected for characterization based on their expression levels, differential expression patterns (more than 4-fold) in young and mature samples, and similarity to characterized CrMDRs. Among the 19 candidates, nine showed higher expression in the mature leaves (*MsMDR*1-9), and 10 showed higher expression in the young leaves (*MsMDR*10-19).

### PPI screening in yeast discovered five novel MsSGD1-MsMDR enzyme complexes

Although both MsSGD1 and MsSGD2 are functional in our assay, the considerably lower activity of MsSGD2 implies a side reaction due to substrate promiscuity, similar to the reported side reaction catalyzed by *R. serpentina* raucaffricine glucosidase^23^. Therefore, we postulated that MsSGD1 is the primary enzyme catalyzing strictosidine deglycosylation in the leaf of *M. speciosa* and used only MsSGD1 as the bait to screen for interacting MsMDRs.

We screened all 19 MsMDR candidates’ interactions with MsSGD1 in yeast to identify novel MsSGD1-MsMDR complexes (**Fig. 4**). For BiFC screening, NV-MsMDR or NV-Ps4’OMT and CV-MsSGD1 were co-expressed in yeast. To identify the localization of each enzyme in yeast, 20 multi-copy plasmids harboring the eCFP-MsSGD1 and 19 eGFP-MsMDR candidates were constructed and expressed in yeast, respectively. Each eGFP-MsMDR construct was also co-expressed with eCFP-MsSGD1 to examine whether the co-expression would re-localize the MsMDR candidate with MsSGD1.

**Fig. 4.**
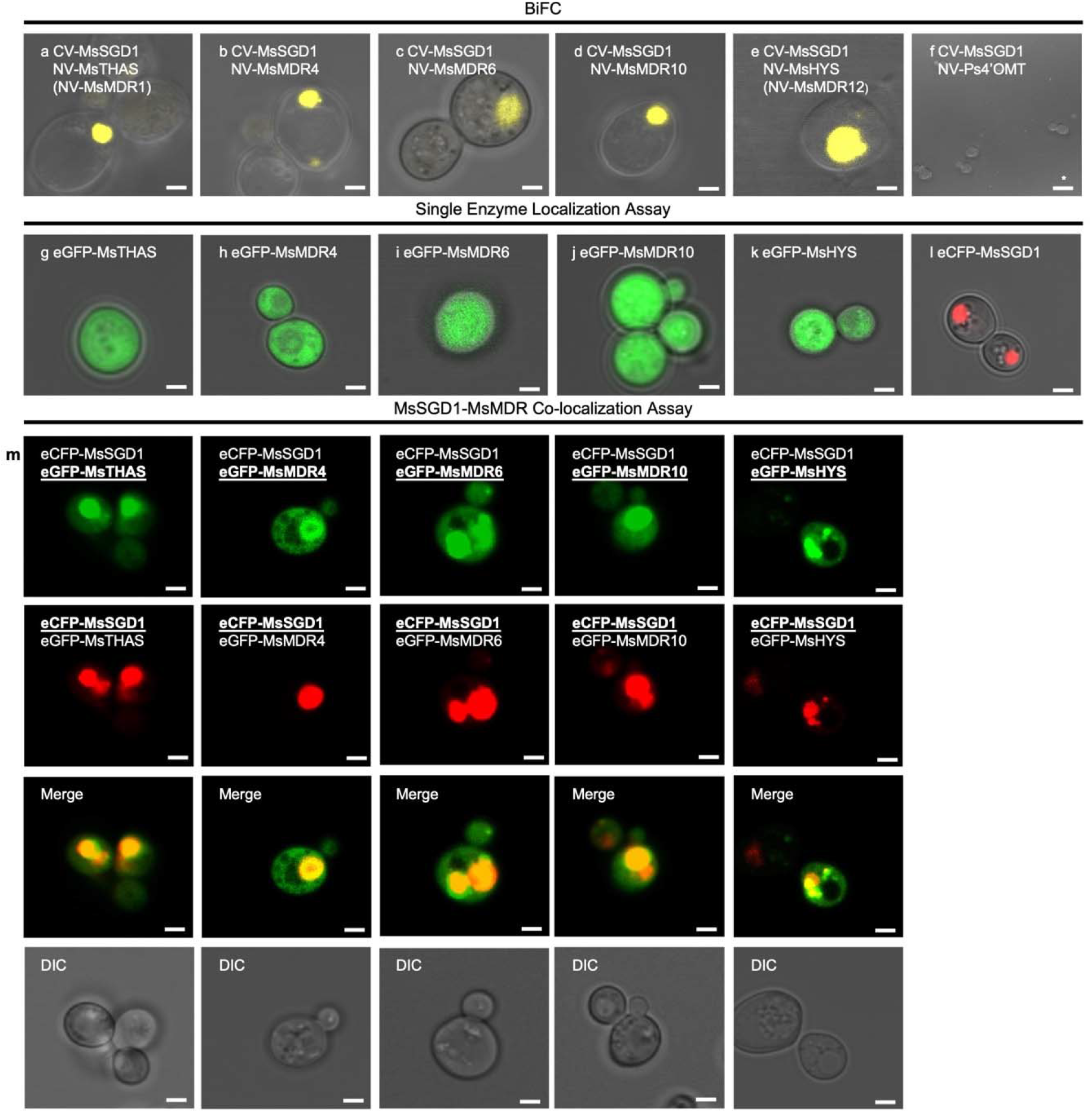
Identified MsSGD1-MsMDR interactions. a-f) MsSGD1-MsMDR interactions were analyzed by BiFC in yeast cells co-expressing CV-MsSGD1 and NV-MsMDRs. MsTHAS (MsMDR1) (a), MsMDR4 (b), MsMDR6 (c), MsMDR10 (d), and MsHYS (MsMDR12) (e) exhibited positive interactions with MsSGD1 in yeast nucleus. Fluorescence signals were shown in yellow false color. The negative control NV-Ps4’OMT (f) did not show fluorescence signal. g-l) Single enzyme localization assay of yeast cells expressing eCFP-MsSGD1 or eGFP-MsMDRs, respectively. MsTHAS (g), MsMDR4 (h), MsMDR6 (i), MsMDR10 (j), and MsHYS (k) localized in yeast cytoplasm. MsSGD1 localized in yeast nucleus (l). Fluorescence signals were shown in red or green false colors. m). Co-localization of co-expressed MsSGD1 and MsMDRs, respectively. MsTHAS and MsMDR10 re-localized in yeast nucleus entirely. MsMDR4, MsMDR6, and MsHYS re-localized in both yeast nucleus and cytoplasm. MsSGD1 localized in yeast nucleus in all six groups. Fluorescence signals were shown in red or green false colors. The co-localization of MsSGD1 and MsMDRs appears in yellow when merging the two individual (red/green) false color images. Cell morphology is observed with differential interference contrast (DIC). All scale bars are 2 μm except for the one labeled with an asterisk (15 μm).

Among the 19 MsMDR candidates screened, five showed positive PPI in both BiFC and co-localization assays and led to novel enzyme complexes in yeast (**Fig. 4**), including MsMDR1 (later named MsTHAS), 4, 6, 10, and MsMDR12 (later named MsHYS). BiFC assays revealed that all PPIs took place in the yeast nucleus (**Figs. 4a-4e**), while the negative control Ps4’OMT did not yield any fluorescence (**Fig. 4f**). Single protein localization assays confirmed that MsSGD1 localized in the yeast nucleus (**Fig. 4l**), and all five MsMDR candidates localized in the cytoplasm (**Figs. 4g-4k**). Co-expressing MsMDRs with MsSGD1 changed the localization of all MsMDR candidates: MsTHAS and MsMDR10 re-localized to the nucleus entirely in the presence of MsSGD1, while MsMDR4, MsMDR6, and MsHYS re-localized in both nucleus and cytoplasm and aggregated preferentially in the nucleus (**Fig. 4m**).

### Biochemical characterization proved three MsSGD1-MsMDR enzyme complexes are functional with MIA products

We then characterized the biochemical activity of the MsMDRs to prove our hypothesis that plant enzymes involved in a complex catalyze sequential reactions in a biosynthetic pathway. 19 engineered yeasts co-expressing MsSGD1 and MsMDRs were used for the crude cell lysate assays with 0.1 mM of crude strictosidine as the substrate. Three out of the five interacting MsMDRs converted strictosidine aglycone to downstream heteroyohimbine-type MIAs or unknown MIA derivatives (**Fig. 5**), including MsHYS, MsTHAS, and MsMDR4. For comparison, we also screened the remaining 14 MsMDR candidates that did not show obvious PPI with MsSGD1, none of which showed detectable products (data not shown).

**Fig. 5.**
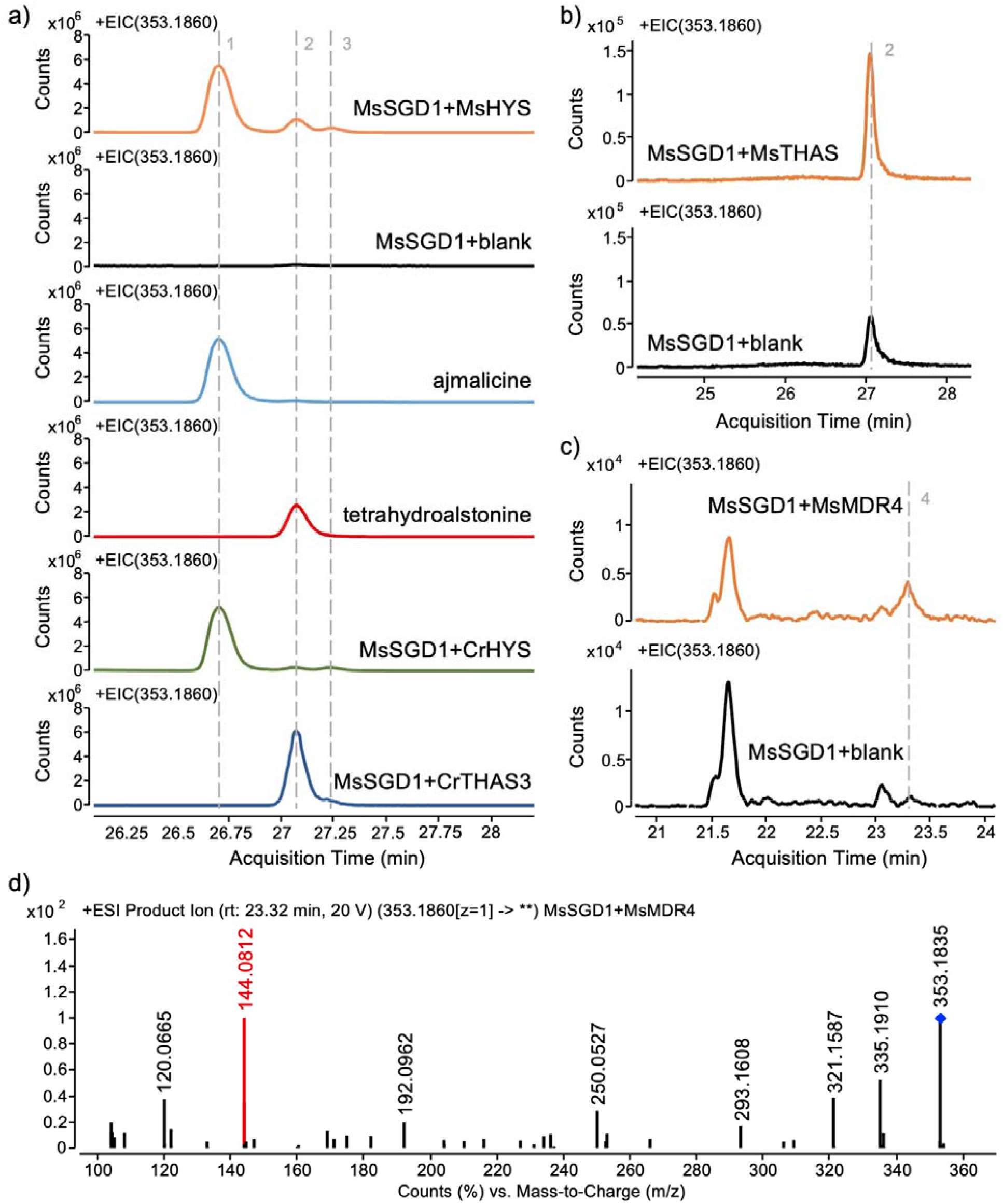
Biochemical characterization of MsHYS, MsTHAS, and MsMDR4. EIC of *m/z* 353.1860 in MsHYS (a), MsTHAS (b), and MsMDR4 (c) characterization experiments. Peak **1, 2**, and **3**: ajmalicine, tetrahydroalstonine, and mayumbine. Peak **4** is an unknown compound, the MS/MS spectrum of which (d) shows a fragment with *m/z* 144.0812, indicating the existence of an indole-related moiety.

MsHYS produced three MS peaks showing the *m/z* of 353.1860 (20 ppm) (**Fig. 5a**). The first two products were identified as ajmalicine and tetrahydroalstonine after comparing their retention times (26.69 min and 27.06 min), MS, and tandem MS/MS spectra with standards. The third product (retention time 27.25 min) was then proven to be mayumbine by expressing CrHYS and *C. roseus* tetrahydroalstonine synthase 3 (CrTHAS3)^14^ and comparing the products (**Fig. 5a**). MsTHAS produced only one product identified as tetrahydroalstonine (**Fig. 5b**). MsMDR4 produced a trace amount of an unknown compound with the *m/z* of 353.1860 (20 ppm) (**Fig. 5c**). MS/MS analysis showed an indole-related moiety (*m/z* of 144.0812, **Fig. 5d**) that was also observed in ajmalicine and tetrahydroalstonine, indicating the production of an MIA. As this product was not present in the leaf extract of *M. speciosa*, it is likely a downstream intermediate from strictosidine aglycones. MsHYS has the highest activity among the three MsMDRs, corresponding with the observation that ajmalicine is the major heteroyohimbine-type MIA synthesized in *M. speciosa*. Sequence analysis revealed that MsHYS shared only 64% amino acid sequence similarity with CrHYS, while MsTHAS shared 58% similarity with CrTHAS1 and 52% with CrTHAS3.

Among the five newly discovered MsSGD1-MsMDR enzyme complexes, three have been proven to produce downstream MIA products, leading to the identification of three functional MsMDR enzymes immediately downstream of MsSGD1. The high agreement between biochemical activities and the PPI screening results proved our hypothesis that the discovery of enzyme complexes can guide novel PNP pathway identification with a much higher success rate than purely transcriptomics analysis. The metabolic functions of MsSGD1-MsMDR6 and MsSGD1-MsMDR10 complexes have not been elucidated. Further optimization of the *in vitro* enzymatic assay would be necessary to determine whether the inability to observe MIA products from these two complexes resulted from the dynamic and unstable nature of the products or from the lack of enzymatic activity. It is also possible that the two-component enzyme complexes are incomplete. It would be helpful to screen for more enzymes (e.g., MsMDRs and cytochrome P450s) that could interact with the MsSGD1-MsMDR scaffold and lead to a complete enzyme complex with catalytic activities.

### Development of heteroyohimbine-type MIA biosynthetic pathway in yeast

We reconstituted the heteroyohimbine-type MIA biosynthetic pathway by co-expressing MsSTR, MsSGD1, and MsHYS in yeast and fed 0.5 mM secologanin and 0.5 mM tryptamine during fermentation. Heteroyohimbine-type MIA was detectable after 24 hours of fermentation (**Figs. 6a** and **6b**). After 72 hours, the engineered yeast produced 1.76 μM ajmalicine, 1.33 μM mayumbine, and 0.40 μM tetrahydroalstonine, respectively (**Fig. 6b**). Approximately 50 μM tryptamine was converted to strictosidine during the fermentation (data not shown). The ratio among the three MIA products (ajmalicine: mayumbine: tetrahydroalstonine=1:0.75:0.09) was different from that obtained in the *in vitro* cell lysate assay (1:0.06:0.16). Although both the *in vivo* and *in vitro* assays were performed in a pH-neutral environment, the *in vivo* fermentation exhibited an obvious product preference for mayumbine over tetrahydroalstonine throughout the 72-hour fermentation process. As the *in vitro* assay lacks the intact cellular context, and the PPI between MsSGD1 and MsHYS has not been validated in the *in vitro* settings, we postulated that the MsSGD1-MsHYS complex was not present or could not regulate the MIA biosynthesis efficiently. Meanwhile, the formation of the MsSGD1-MsHYS enzyme complex in the cellular context might alter the stereospecificity of the reactions via metabolite channeling and re-direct more metabolic flux to mayumbine. It is intriguing that similar changes were observed in the *in vitro* and *in planta* assays of CrTHAS1. When the *in vitro* product of CrTHAS1 was tetrahydroalstonine, CrTHAS1 silencing *in planta* significantly decreased the production of both tetrahydroalstonine and ajmalicine^14^. Taken together, the change of products in both *C. roseus* and *M. speciosa* enzyme complexes implies that SGD-MDR complexes might play an important role in re-directing the metabolic fluxes among various pathway branches.

**Fig. 6.**
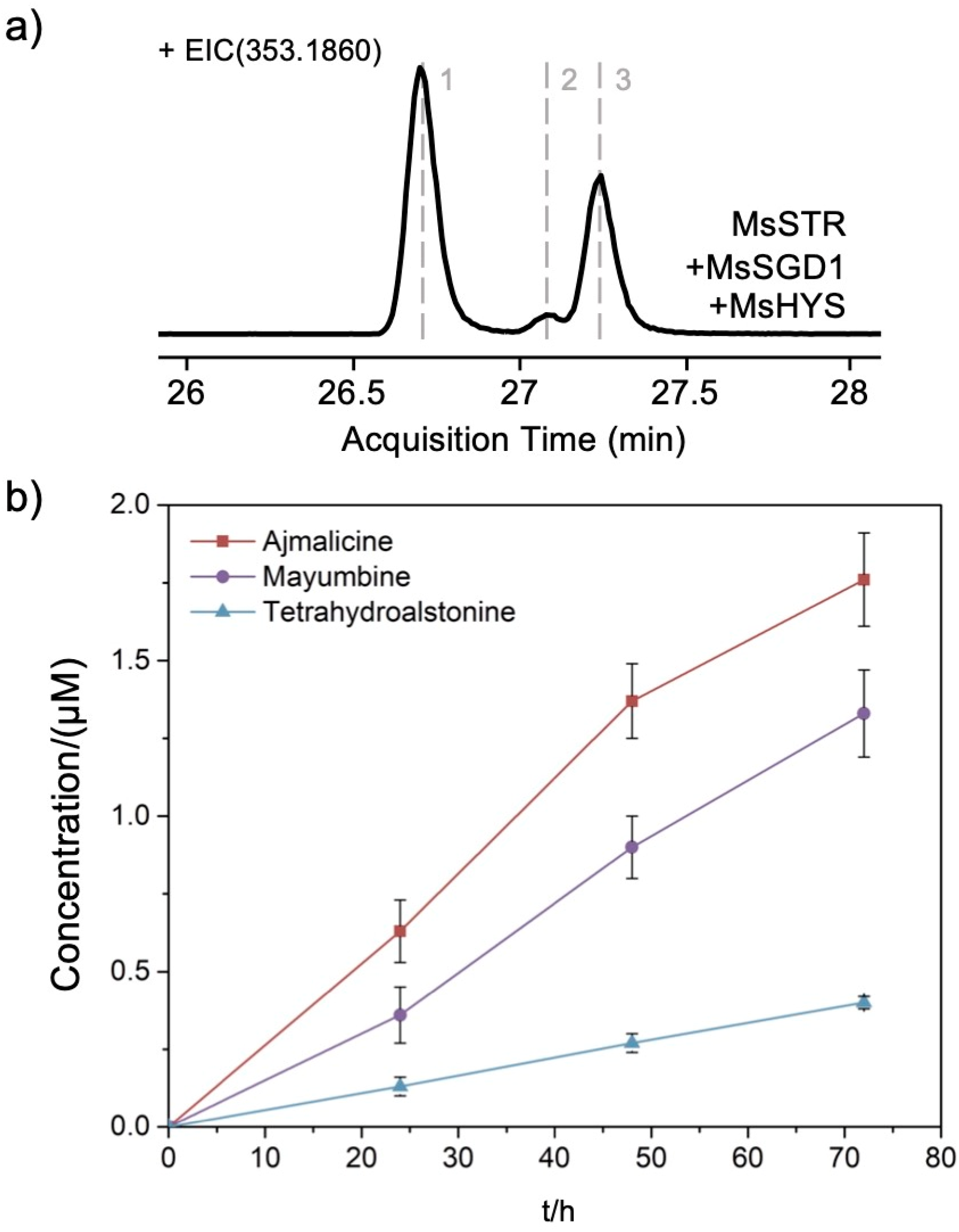
Heteroyohimbine-type MIAs biosynthesis in yeast by reconstructed *M. speciosa* pathway. a) EIC of *m/z* 353.1860 after 24-hour fermentation. Peak **1, 2**, and **3**: ajmalicine, tetrahydroalstonine, and mayumbine. b) Time curves of heteroyohimbine-type MIAs production in 72-hour fermentation. Ajmalicine and tetrahydroalstonine were quantified by corresponding chemical standards. Mayumbine concentrations were estimated according to the ajmalicine standard curve. Error bars represent standard deviations (n□=□3).

## Discussion

Our work demonstrated the discovery of six enzymes (MsSTR, MsSGD1-2, and three MsMDRs) in the MIA biosynthetic pathway from *M. speciosa*. The three pathway branches led to diverse MIA products and can be leveraged to discover downstream enzymes specific to *M. speciosa* in the future. More importantly, the work proved that enzyme complex can be leveraged as a protein-level pattern to discover PNP pathways with a high success rate, which can complement the transcriptomics-driven pathway prediction strategy and accelerate medicinal plant pathway discovery. The discovery of five MsSGD1-MsMDR complexes led to the identification of three functional MsMDRs, which were later confirmed as the only three functional MsMDRs among the 19 putative MsMDRs we characterized. Considering that the 19 putative MsMDRs have been prioritized results from 190 candidates based on their high and differential expression patterns in leaf samples, we concluded that PPI screening is an efficient (60% success rate) method to identify genes that belong to pathways containing enzyme complexes.

Using a known transcript/gene (bait) to identify other transcripts/genes (prey) in the same pathway has proven an effective method to discover PNP pathway genes at the RNA level by gene co-expression analysis^28^ or at the DNA or epigenomic level by clustered gene mining or co-regulation analysis^29,30^. Here, our work highlighted the possibility of a protein-level bait-prey approach to discover PNP pathways spatially organized in the form of dynamic enzyme complexes. As the PPI between interacting enzymes is highly specific and does not require unique substates or cofactors as in biochemical assays, this bait-prey strategy can be further applied for library-based, large-scale, untargeted complex discovery.

This work also proved that SGD-MDR enzyme complexes extensively exist in different MIA-producing plants and provided a synthetic biology approach to discover novel enzyme complexes systematically. It is believed that enzyme complexes that produce similar PNPs are conserved across different plant species. However, no enzyme complexes in the MIA metabolism were reported in plants except for *C. roseus*. Here we have identified two very conserved *M. speciosa* complexes similar to the identified *C. roseus* complexes producing heteroyohimbine-type MIAs, one unique MsSGD1-MsMDR4 complex of which no counterparts were reported in *C. roseus*. Although the MsSGD1-MsMDR6 and the MsSGD1-MsMDR10 complexes have not been functionally characterized, the validated PPIs suggest that they might be the core scaffold of bigger enzyme complexes and lead to novel biosynthetic pathways. In addition, the discovery of enzyme complexes used to be a byproduct of PNP biosynthetic pathway elucidation on an ad hoc basis, after all enzymes had been characterized biochemically. Here we demonstrated that PPI screening could discover novel enzyme complexes from uncharacterized putative enzymes, which has the potential of transforming the current case-by-case discovery process into a more efficient and universal approach.

Key to the efficient discovery of *M. speciosa* enzyme complexes and corresponding MIA pathways is the high-throughput PPI screening in yeast. Unlike *C. roseus*, in which abundant *in-planta* engineering tools and methods have been available, most medicinal plants, including *M. speciosa*, are challenging to engineer directly. The combination of heterologous plant enzyme expression and complementary PPI identification methods enabled us to identify enzyme complexes that used to be inaccessible. However, multiple factors need to be considered before applying the yeast-based PPI screening approach to other plants or complexes. The transient PPIs that lead to the formation of a dynamic plant enzyme complex are fragile, reversible, and usually involve organelle-associated enzymes. Consequently, they are notoriously difficult to validate with high false positive/negative ratios. It is essential to combine multiple PPI assays and to evaluate the results carefully. In our work, we selected BiFC for molecular-level PPI validation as it is compatible with all kinds of PPIs regardless of the proteins’ localization. We also tuned the expression levels of the bait and prey and selected a negative control from *P. somniferum* to avoid the self-assembly of mVenus, which might lead to false positive results. Importantly, the protein localization assay proved an effective method to complement the molecular-level BiFC assay and showed a promising success rate in identifying non-interacting proteins. For example, even with fine-tuned BiFC designs, six MsMDR candidates (MsMDR2, 3, 5, 7, 18, and 19) still showed varied yet positive interactions with MsSGD1 in our BiFC assays (**Supplementary Data**). However, none of the six MsMDR were re-localized or aggregated in the nucleus in the presence of MsSGD1, unlike the obvious re-localization observed in the five interacting MsMDRs (**Supplementary Data**). Comparing their localization and distribution in the single and co-localization assays led to a possible conclusion that the positive BiFC results of these six MsMDRs might result from their natural nuclear localization. MsMDR18 is a natural nuclear protein, and the other five MsMDRs showed varied nucleocytoplasmic localization that did not change in the presence of MsSGD1 (**Supplementary Data**). Therefore, the six candidates were determined as non-interacting MsMDRs. Based on the results of different PPI assays, we speculated that the re-localization and aggregation of the prey enzyme in the presence of the bait is an obvious sign indicating enzyme complex assembly. Future optimization that can re-localize the bait and prey to different organelles in yeast by protein engineering would lead to an effective and universal method for more efficient PPI validation in the future. The feasibility of this method has been supported by the validation of the CrSGD-CrHYS PPI in yeast, which largely benefitted from the re-localization of CrHYS into the cytoplasm. Other optimizations would also be necessary to increase the PPI screening efficiency, including well-defined screening criteria and the development of high-throughput methods beyond individual microscopic analysis. Despite these challenges, the development of the protein-level pathway discovery approach has shown promise and will open new opportunities in PNP pathway discovery.

## Methods

### Chemicals and reagents

Yeast nitrogen base (YNB) and amino acid mixtures were purchased from Sunrise Science Products. Ammonium sulfate, dithiothreitol (DTT), tris(2-carboxyethyl) phosphine hydrochloride (TCEP), secologanin, and tryptamine were purchased from Sigma-Aldrich. Ajmalicine, tetrahydroalstonine, and mitragynine were purchased from Neta Scientific. Dextrose, yeast extract (YE), peptone, Luria-Bertani (LB) broth, LB agar, agar, β-nicotinamide adenine dinucleotide phosphate reduced tetrasodium salt (NADPH), acetonitrile, formic acid, and protease inhibitor (100x) were purchased from Thermo Fisher Scientific. All other chemicals, including antibiotics, were purchased from VWR International.

### Microbes and culture conditions

*E. coli* Top 10, ccdB resistant *E. coli*, yeast CEN.PK2-1D and yeast Strain 4^22^ were used in this work. *E. coli* was used for plasmid construction: Top 10 for normal plasmids and *ccdB* resistant one for plasmids with *ccdB* gene. All *E. coli* strains harboring plasmids were cultivated in LB media or LB plates at 37□ with 50 μg/mL of kanamycin or 100 μg/mL of carbenicillin as appropriate. Yeast CEN.PK2-1D was used for test of enzyme activity and PPI interaction herein. Yeast Strain 4 was cultured for *trCrSTR* cloning. CEN.PK2-1D and Strain 4 were cultured in YPD meduma (1% yeast extract, 2% peptone and 2% dextrose, and additional 2.5% agar for plates) at 30°C, 400 rpm. Yeasts with plasmids were cultivated at 30°C, 400 rpm in appropriate synthetic drop-out (SD) liquid media or plates (0.17% YNB, 0.5% ammonium sulfate, 2% dextrose, and amino acid drop-out mixture, and additional 2.5% agar for plates).

### Plant tissue extract preparation

*M. speciosa plants*, “Rifat Thai” (T) and “Malaysian” (M). strains were incubated in the Guterman greenhouse at Cornell University. The root, mature leaf (T_M and M_M, large and dark green, six-week old), young leaf (T_Y and M_Y, small and light green, two-week old), and flower were harvested, flash frozen in liquid nitrogen, and then extracted. The frozen samples were grounded in a mortar on dry ice with a pestle. The powder was transferred into tubes and frozen at -80□ for 2 hours, followed by lyophilization in a FreeZone benchtop freeze dryer (LABCONON) for three days. The dry weight of samples was measured, and 80% methanol with 0.1% formic acid was added toward 0.1 g dry weight/mL. The mixtures were then subjected to 30 minutes of ultrasound treatment with ultrasonic cleaner (VWR). The sonicated samples were filtered with 0.22 μm polyether sulfone filters and then diluted 20-fold for LC-MS analysis.

### Plant leaf RNA purification, sequencing, and transcriptome analysis

Multiple leaves were grounded to extract RNA from each sample (T_M, T_Y, M_M, and M_Y) following the plant tissue extract preparation protocol to reduce the variations. Total RNA was then extracted using the RNeasy Plant Mini Kit (QIAGEN) according to the manufacturer’s instructions. 2 μg of qualified RNA samples, A_260_/A_280_ ≥ 2.0, A_260_/A_230_ ≥ 2.0, without degradation and contamination, were subjected to Novogene (Beijing) for mRNA library preparation and transcriptome sequencing via NovaSeq 6000 platform with read length of paired-end 150 bp, respectively. The raw data (**Supplementary Table**) had good quality and was further trimmed using Trimmomatic^32^ to remove reads containing adapters or with low quality. The clean reads were assembled into T, M, and T&M transcriptomes using Trinity^33^ (**Supplementary Table**), resulting in 115926, 111502, and 138629 trinity ‘genes’, and 215215, 202523, and 262783 trinity transcripts, respectively. The number indicated that around 80% of transcripts were shared in both strains. T&M transcriptome was evaluated as 99.2% complete by the metric of Benchmarking Universal Single-Copy Orthologs^34^ and was used for downstream analysis. The 129489 putative complete or incomplete open reading frames predicted from Transdecoder were annotated with Trinotate^35^ for gene mining. Gene expression levels were estimated by RSEM^36^. Differential expression analysis was performed using edgeR^37^ with the dispersion parameter of 0.1 and *p*-value of 0.001.

### Plant cDNA preparation

Plant cDNA was prepared from 3 μg of each leaf RNA sample via RNA to cDNA EcoDry Premix (Takara Bio) according to the manufacturer’s instructions. After reverse transcription, the product was cleaned with DNA clean kit (Zymo Research) to get around 180 ng of cDNA for each sample.

### Plasmid construction for gene expression in yeast

Reported monoterpene indole alkaloids biosynthetic genes, *CrSGD, CrHYS*, and *CrTHAS3* from *C. roseus*, and *RsSGD* from *R. serpentina* were codon-optimized and synthesized by Twist Bioscience. *trCrSTR* was amplified from yeast Strain 4 genomic DNA. Other enzyme-encoding genes were amplified from plant cDNA. Gene sequences were listed in **Supplementary Table**. Oligonucleotide primers (**Supplementary Table**) were synthesized by Life Technologies. Enzyme-encoding genes were inserted into pre-assembled Gateway-compatible plasmids that contain constitutive promoters and terminators (**Supplementary Table**). The gene expression cassettes were assembled into multi-copy expression plasmids, including pAG423-ccdB, pAG424-ccdB, and pAG4U6-ccdB, using Gateway LR Clonase II Enzyme mix (Life Technologies), respectively. Plasmids were extracted using plasmid miniprep kits (Zymo Research), and the sequences were confirmed by Sanger Sequencing (Biotechnology Resource Center, Cornell University).

### *In vivo* assays of CrSTR and MsSTR using engineered yeast

Plasmids harboring plant genes were transformed into *S. cerevisiae* CEN.PK2-1D using frozen-EZ yeast transformation II kits (Zymo Research). The resulting yeast strains were cultivated at 30°C, 400 rpm in appropriate SD liquid media or plates and tested as the following in triplicates.

CEN.PK2-1D strains containing pAG424-ccdB, pAG424-CrSTR, or pAG424-MsSTR were cultivated in 0.5 mL of SD-Trp media for two days in a 96-well deep well plate. The seed cultures were then back-diluted 20-fold into fresh SD-Trp media for another two days of incubation. The cultures were centrifuged in the 96-well plate at 4000 rpm. The pellets were washed with fresh SD-Trp media, centrifuged, and resuspended in feeding media (SD-Trp media with 0.5 mM secologanin, 0.5 mM tryptamine, 100 mM Tris-Base, pH7.5, filtered) for 48 hours of reaction in the shaker. After adding 0.1 μM of berberine as the internal standards, samples were harvested, and the supernatants were used for LC-MS analysis after centrifugation at 15000 rpm for 10 minutes.

### *In vitro* assays of plant enzymes with crude yeast cell lysate

CEN.PK2-1D containing pAG426-ccdB, pAG426-CrSGD, pAG426-RsSGD, or pAG426-MsSGDs were cultivated in 1 mL of SD-Ura media in 14-mL culture tubes. CEN.PK2-1D co-expressing pAG4U6-MsMDRs with pAG423-MsSGD1, respectively, were cultured in 1 mL of SD-His-Ura media. The seed cultures were back-diluted 100-fold into 5 mL of fresh media in 14-mL culture tubes. After one day, the cell density was measured, and cultures were diluted to OD 5. 5 mL of diluted cultures were centrifuged at 4000 rpm, 4□ for 5 min. The pellets were resuspended with 250 μL of Tris-HCl breaking buffer (10 mM Tris-HCl, 1% glycerol, 0.1 mM TCEP, 1x protease inhibitor, pH=7.4) after removing the supernatants. The resulting cultures were centrifuged at 10000 rpm, 4□ for 1 min, and the pellets were resuspended with 25 μL of Tris-HCl breaking buffer with 4 mM DTT after removing the supernatants. The resuspended cultures were lysed in microtubes with 425-600 μm glass beads (Sigma-Aldrich) using a BeadBug6 microtube homogenizer (Benchmark Scientific) for twelve bursts of 30 seconds with a 30-second rest in between cycles. 30 μL of Tris-HCl breaking buffer was added into the lysed cell lysate, and 25 μL of the mixtures were harvested for cell lysate assays with 1 μL of crude strictosidine (around 2.5 mM, prepared from the fermentation supernatant of trCrSTR assay by lyophilization, concentration in 80% methanol, and centrifugation to remove the pellets) for the functional characterization of SGD, and an addition of 0.5 μL of 250 mM NADPH for the characterization of MDR. The reaction was then performed at 30□, 400 rpm for one hour. Three volumes of methanol with 0.13% formic acid were then added into the tubes to stop the reaction. After centrifugation at 15000 rpm for 15 minutes, the supernatants were used for LC-MS analysis.

### Characterization of SGD-MDR interaction by microscopy-based bimolecular fluorescence complementation (BiFC) assay

The BiFC assay was developed based on the split-mVenus fluorescent protein. Genes encoding SGD, MDR, and negative control *Pm4’OMT* (synthesized by Twist Bioscience) were cloned into the Gateway pENTR plasmid using Gibson Assembly and recombined using the Gateway LR Clonase II Enzyme mix (Life Technologies). Resultant plasmids were co-transformed into CEN.PK2-1D using the frozen-EZ yeast transformation II kits (Zymo Research). Corresponding yeast strains were cultivated at 30□, 400 rpm in SD-Trp-Leu for three days. Cultured cells were washed with the PBS buffer (137 mM NaCl, 2.7 mM KCl, 10 mM Na_2_HPO_4_, and 1.8 mM KH_2_PO_4_) three times and resuspended with equ-volume PBS. Microscopic analysis was performed using a Zeiss LSM 710 Confocal Microscope (AxioObserver, with objective Plan-Apochromat 63X/1.40 Oil DIC M27) under the LineSequential scanning mode, with the excitation wavelength of 514 nm and signals of 519-620 nm. Transmitted light images (bright field and DIC) were also recorded.

### Characterization of subcellular localization and analysis of co-localizations under confocal microscopy

To fuse the SGD and MDR enzymes with eCFP and eGFP fluorescence proteins, the pENTR plasmids constructed were used for Gateway cloning. The genes were subsequently recombined into expression vectors from the *S. cerevisiae* Advanced Gateway Destination Vector Kit^38^ using LR Clonase II to generate the corresponding yeast expression constructs. Plasmids expressing eCFP-SGD and eGFP-MDR were co-transformed into CEN.PK2-1D using the frozen-EZ yeast transformation II kits (Zymo Research). Resultant yeast strains were cultivated at 30 □, 400 rpm in SD-Trp-Ura media for 16-18 hours and prepared as demonstrated above. Microscopic analysis was performed using a Zeiss LSM 710 Confocal Microscope (AxioObserver, with objective Plan-Apochromat 63X/1.40 Oil DIC M27). For the detection of eCFP fluorescence, the excitation wavelength was 405 nm, and signals of 442-510 nm were recorded by a PMT detector. For the detection of eGFP fluorescence, the excitation wavelength was 488 nm and signals of 491-588 nm were recorded by a PMT detector. Transmitted light images (bright field and DIC) were also recorded.

### Heteroyohimbine biosynthesis in yeast

Plasmids were co-transformed into CEN.PK2-1D using the frozen-EZ yeast transformation II kits (Zymo Research). The resulting strains were then tested by secologanin and tryptamine feeding in appropriate media using the same protocol as *in vivo* assays of CrSTR and MsSTR. Products in the strain cultures were analyzed 24-, 48- and 72-hour post-feeding by LC-MS.

### LC-MS analysis

Metabolites were analyzed by an HPLC-Q-TOF (Agilent 1260 Infinity II/Agilent G6545B) in MS mode using positive ionization, with water with 0.1% formic acid (A) and acetonitrile with 0.1% formic acid (B) as the mobile phase. For STR assays, 1 μL of the sample was injected and separated in the Agilent ZORBAX RRHD Eclipse Plus C18 column (2.1 × 50 mm, 1.8 μm) with a short gradient program (0-1 min, 95% A; 1-11 min, 95%-5% A; 11-13 min, 5% A; 13-14 min, 5%-95% A; and 14-16 min, 95% A) at 40□ and a flow rate of 0.4 mL/min. For efficient separation of compounds in tissue extract, the short gradient program was extended to 60 minutes (0-4 min, 95% A; 4-36 min, 95%-60% A; 36-45 min, 60%-5% A; 45-52 min, 5% A; 52-56 min, 5%-95% A; and 14-16 min, 95% A) instead. The sample injection volume was further increased to 4 μL for product analysis in SGD and SGD+MDR cell lysate assays. Specifically, to distinguish between ajmalicine and tetrahydroalstonine isomers, samples were injected and separated in a long Agilent ZORBAX RRHT Eclipse Plus C18 column (4.6 × 150 mm, 1.8 μm) with the 60-min elution method at 60□ and a flow rate of 0.6 mL/min. The *m/z* value of the [M+H]^+^ adduct was then used to extract the ion chromatogram (with a mass error below 20 ppm) for compound identification or quantification with corresponding chemical standards. For further identification of compounds, target MS/MS mode with 20 eV of collision energy was used in addition to high-resolution MS analysis.

### Data Availability

RNA Sequencing data have been deposited into the National Center for Biotechnology Information, NIH, Sequence Read Archive (accession xxxxxxxxxxx). The sequences of the functional genes reported in this article have been deposited in NCBI GenBank (accessions xxxxxxxx-xxxxxxxx). All other data supporting the findings of this study are presented in the published article (including its Supplementary Information) or are available from the corresponding author upon reasonable request.

## Acknowledgements

This work was supported by the National Institutes of Health - National Institute on Deafness and Other Communication Disorders under award number R21DC019206 to S. Li, the National Institutes of Health - National Institute of General Medical Sciences under award number T32GM138826, the National Science Foundation under Grant No. DBI-2019674, and the Research Innovation Fund from Cornell Institute for Digital Agriculture. We thank the Bioinformatics Facility (RRID: SCR_021757) and the Imaging Facility (RRID: SCR_021741) of the Biotechnology Resource Center of Cornell Institute of Biotechnology. We thank S. E. O’Connor from Max Planck Institute of Chemical Ecology for kindly sharing Strain4. We thank J. Han for help in developing PPI assays. We thank D. Han, J. Han, Y. Huang, and A. Koganitsky for their valuable comments on this manuscript.

## Author contributions

Y.W., C.L., and S.L. designed the experiments; Y.W., C.L., and F. G. performed experiments; Y.W., C.L., and S.L. analyzed the data and wrote the manuscript.

